# Molecular and cellular characterization of the interaction between the Rho GAP RGA-3/4 and F-actin

**DOI:** 10.64898/2026.09.18.752697

**Authors:** Songeun Kim, Kevin J. Sonnemann, Andrea A. Putnam, William M. Bement

## Abstract

During cytokinesis in amphibian and echinoderm embryos, propagating waves of Rho activity and F-actin assembly – cortical excitability – are amplified and focused at the equatorial cortex by the mitotic spindle. Such cortical waves are proposed to result from fast positive feedback based on Rho and the Rho GEF, Ect2, and delayed negative feedback involving F-actin and the Rho GAP, RGA-3/4. Currently, this model is tenuous, in part because the relationship between F-actin and RGA-3/4 is not well understood. Therefore, we investigated how RGA-3/4 regulates F-actin structure both *in vivo* and *in vitro*. Full-length RGA-3/4, when expressed alone in *Xenopus* oocytes, localizes to and bundles cortical F-actin. *In vitro*, purified full-length RGA-3/4 protein directly binds to and bundles F-actin. TIRF microscopy revealed that RGA-3/4 binds to F-actin cooperatively and accumulates on the filaments before filament crosslinking events. Structure-function analyses demonstrated both the amino-terminal region and the intrinsically disordered region at the carboxyl-terminal portion of RGA-3/4 are essential for normal F-actin interaction. These results provide the first demonstration that RGA-3/4 is a direct, cooperative F-actin binding and bundling protein. We propose that its special features make RGA-3/4 uniquely suited to provide F-actin-dependent negative feedback during cortical excitability and cytokinesis.

**Significance statement:**

- Cortical excitability during cytokinesis relies on delayed negative feedback based on F-actin and the Rho GAP, RGA-3/4. However, the molecular basis of the interaction between F-actin and RGA-3/4 is uncharacterized.
- We show that RGA-3/4 directly and cooperatively binds to and bundles F-actin and identify the regions of RGA-3/4 required for the F-actin interaction.
- Our work provides a new framework to understand how RGA-3/4 contributes to actin remodeling in cytokinesis and other essential morphogenetic processes.

## Introduction

The cell cortex – the plasma membrane and the underlying actomyosin cytoskeleton – is a dynamic interface where biochemical signaling and cytoskeletal organization are coordinated to generate diverse patterns (Bement *et al*., 2006). One fascinating type of cortical pattern is cortical excitability, a behavior characterized by pulses or propagating waves of F-actin assembly and the activity of actin regulators, including Rho GTPases and their binding partners (Michaud *et al*., 2021). Cortical excitability is essential for various morphogenetic events that require precise spatiotemporal control of cell shape, contractility, and force production, including cell migration, cell division, embryo compaction, exocytosis, adhesion, polarization, and germline cell sorting (Vicker, 2000; Bement *et al*., 2015; Maître *et al*., 2015; Rousso *et al*., 2016; Graessl *et al*., 2017; Xiao *et al*., 2017; Michaux *et al*., 2018; Chanet and Huynh, 2020; Michaud *et al*., 2022).

In frog and starfish embryos, cortical excitability is manifested as propagating waves of Rho GTPase activity and complementary waves of F-actin assembly that are amplified and focused at the equatorial cortex by the mitotic spindle, where they are responsible for the assembly of the cytokinetic apparatus which constricts the cell in half (Bement *et al*., 2015; Michaud *et al*., 2022; Swider *et al*., 2022). These wave dynamics are proposed to result from coupling of two feedback loops: fast positive feedback loop dependent on Rho and the Rho GEF, Ect2 and delayed negative feedback loop involving F-actin and the Rho GAP, RGA-3/4 (Michaud *et al*., 2022).

RGA-3/4, also known as ArhGAP11A and MP-GAP (Zanin *et al*., 2013), is a Rho GAP whose homologs have been implicated in diverse biological contexts including cell polarity, cytokinesis, cortical excitability, and mammalian brain development (Schonegg *et al*., 2007; Zanin *et al*., 2013; Michaux *et al*., 2018; Michaud *et al*., 2022, Hass *et al*., 2025). RGA-3/4 has also attracted considerable interest in human disease as both a potential prognostic biomarker and a therapeutic target in multiple cancers, including breast, lung, gastric, renal, and pancreatic cancer (Lawson *et al*., 2016; Chen *et al*., 2021; Fan *et al*., 2021; Yang *et al*., 2023; Shu *et al*., 2025). Despite its importance, we still know surprisingly little about how RGA-3/4 functions at a molecular level. Previous work has shown that cortical recruitment of RGA-3/4 depends on F-actin (Michaux *et al*., 2018; Michaud *et al*., 2022). However, the molecular basis of RGA-3/4 interaction with F-actin remains poorly understood. One challenge is that RGA-3/4 has primarily been studied in dynamic cortical processes, where its recruitment occurs in complex spatially- and temporally-patterned behaviors. In these processes, RGA-3/4 recruitment coincides with F-actin recruitment and the recruitment of other actin-associated proteins (Michaux *et al*., 2018; Michaud *et al*., 2022), making it difficult to determine whether RGA-3/4 is directly recruited to F-actin itself or to an F-actin binding protein. This uncertainty is exacerbated by the fact that RGA-3/4 lacks any canonical F-actin binding motifs or other well-defined non-catalytic regions. Moreover, because full-length RGA-3/4 has never been purified, the possibility that it binds to F-actin by non-canonical means has not been assessed.

Here, we investigated the interaction between RGA-3/4 and F-actin in immature *Xenopus*oocytes and *in vitro*. Immature *Xenopus* oocytes are arrested in meiotic interphase and are quiescent in terms of cortical waves and contractility since the main regulators of excitability including RGA-3/4 are not expressed at this cell stage. Thus, the *Xenopus* oocytes provide a stable *in vivo* platform wherein we could directly test whether and how RGA-3/4 interacts with cortical F-actin independently from the context of dynamic Rho-dependent feedback. We also developed the means to purify recombinant full-length RGA-3/4 as well as several mutants which allowed us to directly assess RGA-3/4-F-actin binding *in vitro*. Together, these approaches provide the first direct characterization of RGA-3/4 interaction with F-actin.

### RGA-3/4 bundles and colocalizes with cortical F-actin in immature *Xenopus* oocytes

RGA-3/4 contains a conserved GAP domain that is specific for Rho (Zanin *et al*., 2013) and amino- and carboxyl-terminal regions of unknown function (Figure 1A). Although these presumptively non-catalytic regions are not strongly conserved at the primary sequence level across the homologs, AlphaFold structure prediction suggests that much of the protein outside of the GAP domain is intrinsically disordered (Figure 1A’). To test whether RGA-3/4 can associate with the cortex and colocalize with cortical F-actin in resting cells (i.e. independent of cortical excitability), RGA-3/4-GFP was expressed in immature *Xenopus* oocytes. RGA-3/4-GFP colocalized with cortical F-actin and reorganized it into large bundles (Figure 1B). This phenotype was observed for both *Xenopus* RGA-3/4 and its human homolog (Figure 1B), showing that the ability to bundle cortical F-actin *in vivo* is conserved.

**Figure 1.**
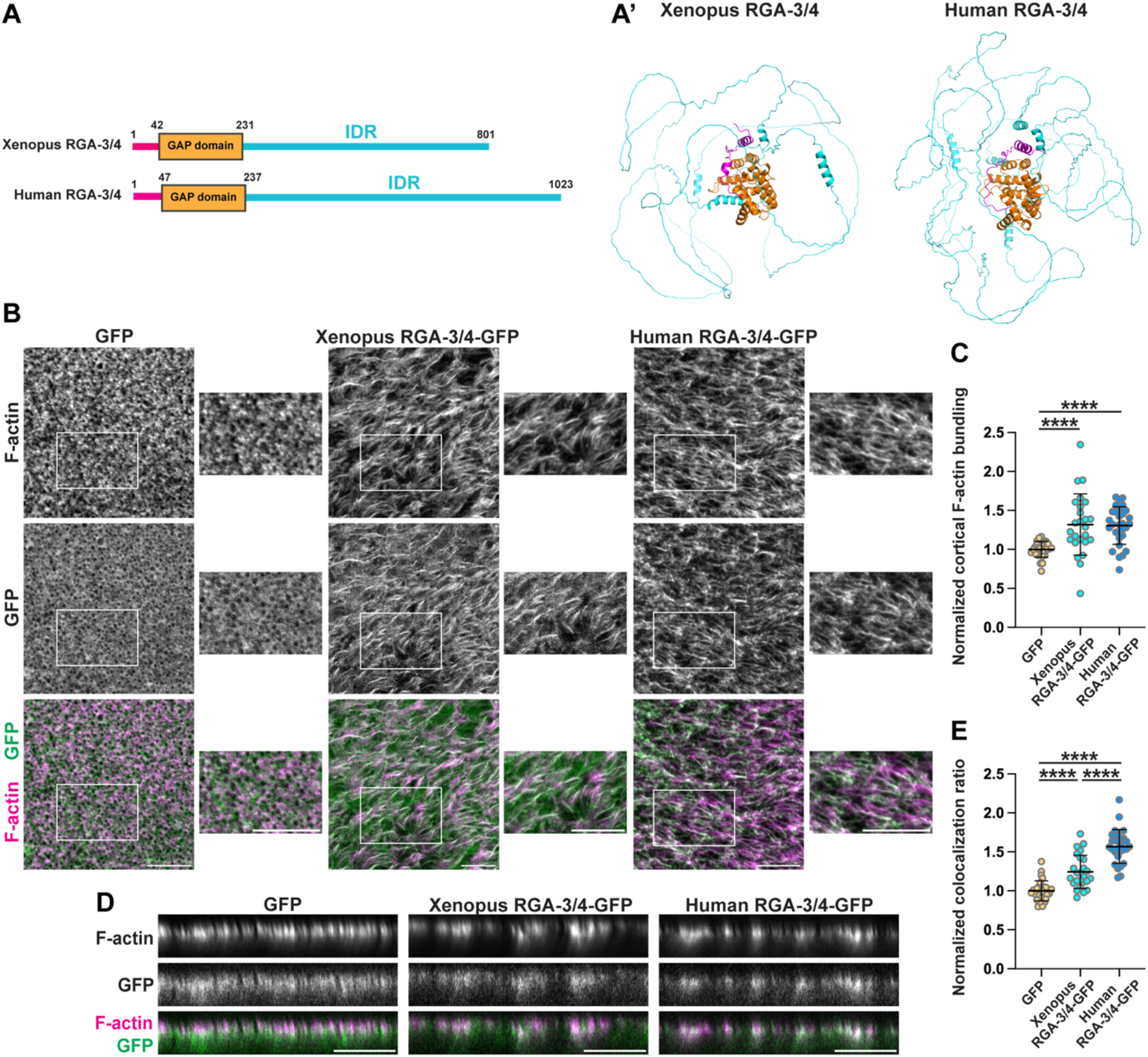
RGA-3/4 bundles and colocalizes with cortical F-actin in immature Xenopus oocytes. (A) Schematic diagrams of Xenopus and Human RGA-3/4 and (A’) the predicted AlphaFold structures. IDR: Intrinsically disordered region. (B) Cortex of Xenopus oocytes expressing GFP (control), Xenopus RGA-3/4-GFP, or Human RGA-3/4-GFP. The oocytes were fixed with Alexa546-phalloidin. Scale bar: 20µm. (C) Quantification of normalized cortical F-actin bundling. GFP, n=29; Xenopus RGA-3/4-GFP, n=25; Human RGA-3/4-GFP, n=30; 3 experiments. ****, <0.0001. (D) Sideview profile of the cortex across experimental groups described in B. Scale bar: 20µm. (E) Quantification of normalized colocalization ratio. GFP, n=29; Xenopus RGA-3/4-GFP, n=25; Human RGA-3/4-GFP, n=28; 3 experiments. ****, <0.0001. (C and E) Mean±SD is shown. Each dot represents measurement in a single oocyte. One-way ANOVA with Tukey post-hoc test for multiple comparisons was performed for statistical analysis.

Because bundled F-actin contains regions with different filament densities, it shows higher variation in signal intensity than control cortical F-actin which is relatively uniform. Cortical F-actin bundling was therefore quantified by using the coefficient of variation of F-actin fluorescence intensity, calculated as the standard deviation divided by the mean intensity. Compared to GFP alone, both RGA-3/4-GFP and its human homolog significantly increased cortical F-actin bundling (Figure 1C). To eliminate the possibility that GFP contributed to the bundling phenotype, we tested untagged RGA-3/4 and found that both untagged *Xenopus* RGA-3/4 and its human homolog induced cortical F-actin bundling (Supplemental Figure S1). We next sought to assess the colocalization of RGA-3/4-GFP with cortical F-actin. Since the signal from free GFP was distributed broadly throughout the cortex and underlying cytoplasm, cortical enrichment of RGA-3/4-GFP was assessed using sideview intensity profiles (Figure 1D). Colocalization was measured as the fraction of GFP signal colocalized with cortical F-actin relative to the total GFP signal detected at the cortex and in the cytoplasm. This analysis showed that RGA-3/4-GFP and its human homolog were significantly enriched at the cortex compared to GFP alone (Figure 1E). Collectively, these results show that RGA-3/4 associates with cortical F-actin and identify its conserved ability to remodel F-actin organization *in vivo*.

### Structure-function analysis of RGA-3/4 interaction with cortical F-actin

Colocalization and bundling were next used as readouts of cortical F-actin interaction in a structure-function analysis of different mutants shown schematically in Figure 2A and 3A. Full-length RGA-3/4 was used as a positive control while free GFP was used as a negative control. All RGA-3/4 constructs were tagged with GFP, allowing confirmation that similar levels of expression were achieved with the various mutants. The isolated GAP domain of RGA-3/4 neither bundled nor colocalized with F-actin (Figure 2, B-D). To determine whether GAP activity or interaction of the GAP domain with Rho contributes to cortical F-actin interaction, two GAP-dead mutants were tested: R80A and R80E. Both mutations are predicted to disrupt the catalytic activity of the GAP domain but they differ in their expected effects on GAP-Rho interaction: R80A is predicted to impair catalytic activity without completely disrupting GAP-Rho complex formation while R80E is expected to abolish GAP-Rho binding (Dvorsky and Ahmadian, 2004). The R80A mutant of RGA-3/4 retained the ability to bundle and colocalize with cortical F-actin, while the R80E mutant failed to bundle or colocalize with F-actin (Figure 2, B-D). These results suggest that rather than catalytic GAP activity itself, the engagement with Rho might contribute to RGA-3/4 interaction with cortical F-actin.

**Figure 2.**
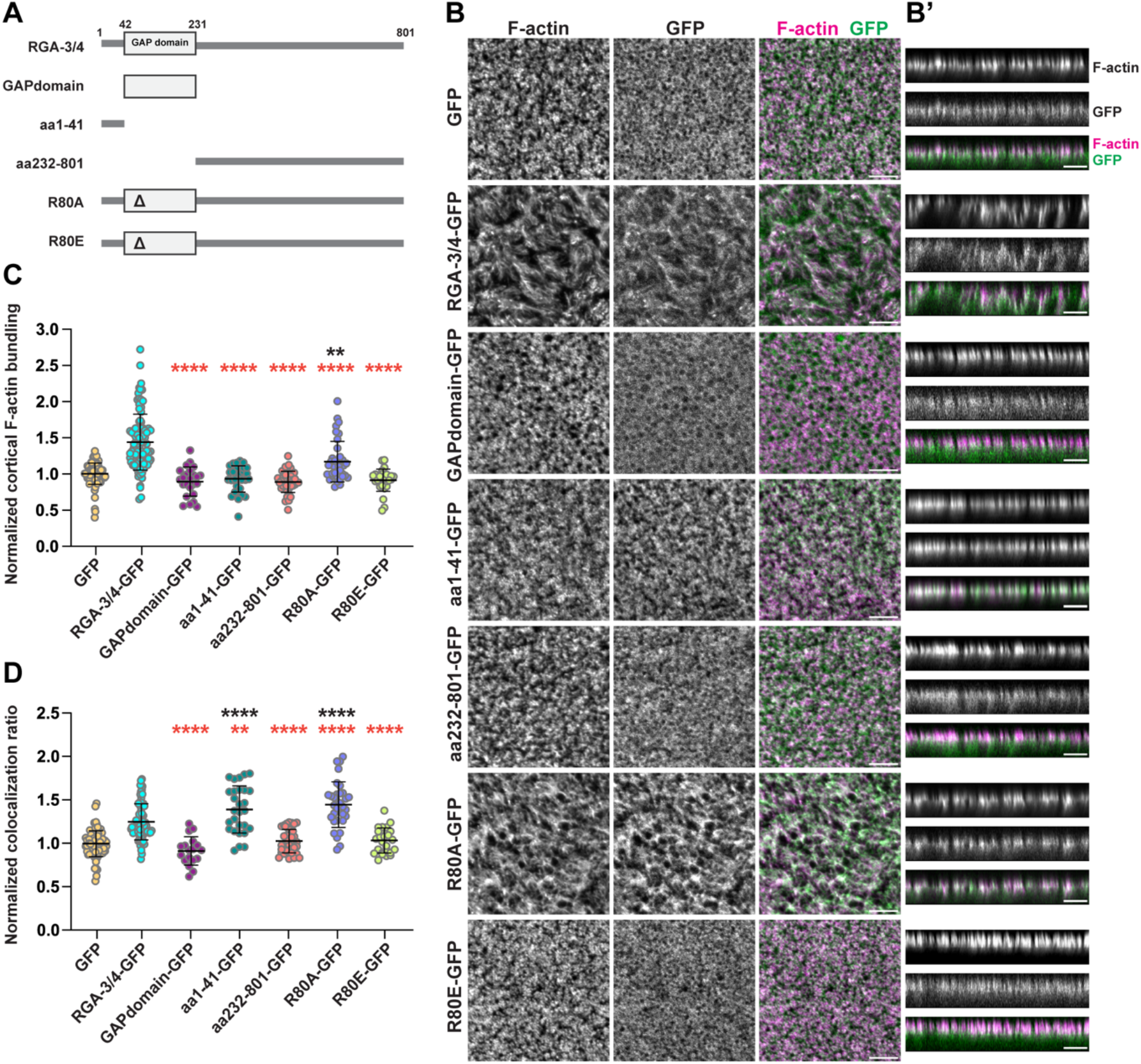
Structure-function analysis of RGA-3/4 interaction with F-actin in Xenopus oocytes. (A) Schematic diagrams of RGA-3/4 and various truncated or mutated constructs. (B) Representative oocyte from each treatment group. En-face and (B’) sideview profiles of the cortex are shown. The oocytes were fixed with Alexa546-Phalloidin. Scale bar: 10*μ*m. (C) Quantification of normalized cortical F-actin bundling. GFP, n=99; RGA-3/4-GFP, n=91; GAPdomain-GFP, n=24; aa1-41-GFP, n=32; aa232-801-GFP, n=42; R80A-GFP, n=38; R80E-GFP, n=29; 13 experiments. **=0.0036; ****, <0.0001. (D) Quantification of normalized colocalization ratio. GFP, n=81; RGA-3/4-GFP, n=71; GAPdomain-GFP, n=18; aa1-41-GFP, n=28; aa232-801-GFP, n=39; R80A-GFP, n=33; R80E-GFP, n=26; 9 experiments. **=0.0065; ****, <0.0001. (C and D) Mean±SD is shown. Each dot represents measurement in a single oocyte. One-way ANOVA with Dunnett post-hoc test for multiple comparisons was performed for statistical analysis. Black asterisks: comparison to GFP (negative control). Red asterisks: comparison to RGA-3/4-GFP (positive control).

**Figure 3.**
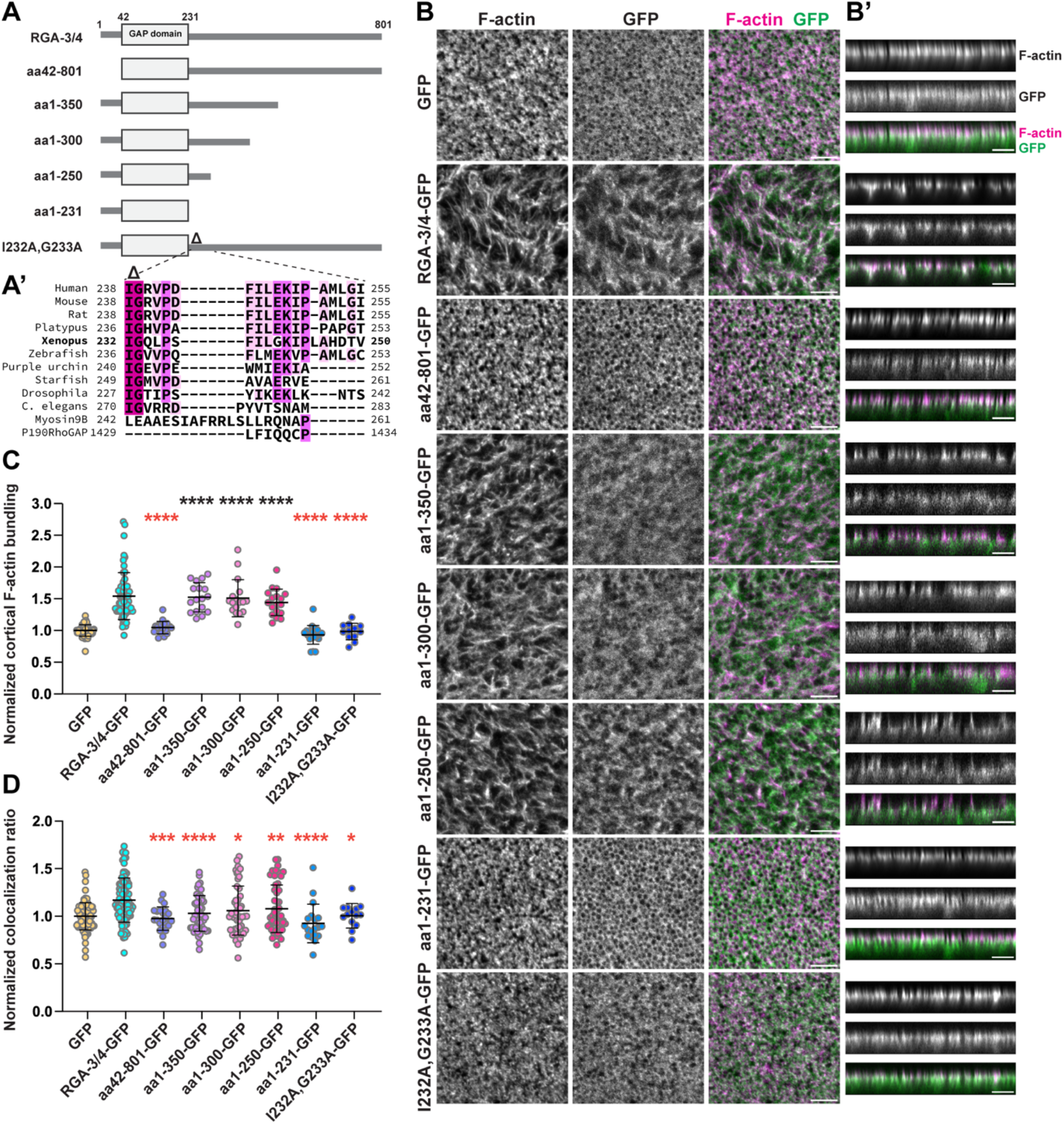
Non-catalytic regions of RGA-3/4 are important for cortical F-actin interaction. (A) Schematic diagrams of RGA-3/4 and various truncated or mutated constructs. (A’) Sequence alignment of various homologs of RGA-3/4 and other Rho GAPs, myosin 9B and P190RhoGAP, for region just downstream of GAP domain. MAFFT sequence alignment program was used. Highlighted residues represent agreement with the alignment’s consensus sequence, with darker shade indicating higher agreement. (B) Representative oocyte from each treatment group. En-face and (B’) sideview profiles of the cortex are shown. The oocytes were fixed with Alexa546-Phalloidin. Scale bar: 10*μ*m. (C) Quantification of normalized cortical F-actin bundling. GFP, n=53; RGA-3/4-GFP, n=60; aa42-801-GFP, n=19; aa1-350-GFP, n=17; aa1-300-GFP, n=17; aa1-250-GFP, n=18; aa1-231-GFP, n=21; I232A,G233A-GFP, n=12; 10 experiments. ****, <0.0001. (D) Quantification of normalized colocalization ratio. GFP, n=99; RGA-3/4-GFP, n=89; aa42-801-GFP, n=22; aa1-350-GFP, n=47; aa1-300-GFP, n=49; aa1-250-GFP, n=51; aa1-231-GFP, n=19; I232A,G233A-GFP, n=14; 14 experiments. *=0.0147 (aa1-300-GFP); *=0.0312 (I232A,G233A-GFP); **=0.0081; ***=0.0004; ****, <0.0001. (C and D) Mean±SD is shown. Each dot represents measurement in a single oocyte. One-way ANOVA with Dunnett post-hoc test for multiple comparisons was performed for statistical analysis. Black asterisks: comparison to GFP (negative control). Red asterisks: comparison to RGA-3/4-GFP (positive control).

We next tested whether regions outside the GAP domain were sufficient for cortical F-actin interaction. The amino-terminal region (aa1-41) did not induce any cortical F-actin bundling but did colocalize with cortical F-actin (Figure 2, B-D). In contrast, the carboxyl-terminal region (aa232-801; contains all of the intrinsically disordered region (IDR) carboxyl-terminal to the GAP domain) both failed to bundle and colocalize with cortical F-actin (Figure 2, B-D). The colocalization of aa1-41 with F-actin suggested that this region may contain an F-actin interaction site. Supporting this idea, a mutant with this region removed (aa42-801) neither bundled nor colocalized with cortical F-actin (Figure 3, B-D). The corresponding human mutant (aa47-1023) also failed to induce cortical F-actin bundling and had reduced colocalization relative to full-length human RGA-3/4 (Supplemental Figure S2).

While the above results indicated the aa1-41 is necessary for F-actin binding and bundling *in vivo*, they also indicated that aa1-41 is not sufficient for F-actin bundling. We therefore tested series of carboxyl-terminal truncations in which the IDR was progressively shortened: aa1-350, aa1-300, aa1-250, and aa1-231. The first three truncated constructs induced cortical F-actin bundling but failed to show colocalization significantly above that seen with GFP alone (Figure 3, B-D). In contrast, aa1-231 failed to either bundle or colocalize with cortical F-actin (Figure 3, B-D). Since aa1-250 induced F-actin bundling while aa1-231 did not, the region in between (aa232-250) may contain an additional F-actin interaction site needed for F-actin bundling. To further examine this region, we aligned RGA-3/4 homologs and found that IG residues located immediately downstream of the GAP domain are highly conserved (Figure 3A’). Notably, these residues appear to be unique to RGA-3/4 as the GAP domains from other Rho GAPs including p190GAP and myosin9B do not contain these residues downstream of their GAP domains (Figure 3A’). Mutation of the conserved IG residues to alanine (I232A,G233A) within full-length RGA-3/4 abolished cortical F-actin bundling and colocalization, confirming the importance of these residues (Figure 3, B-D). The corresponding mutation in the human homolog (I238A,G239A) also significantly reduced the extent of cortical F-actin bundling and colocalization (Supplemental Figure S2). Collectively, the structure-function experiments reveal that multiple parts of RGA-3/4 contribute to its cortical F-actin interaction *in vivo*. That is, the amino-terminal region (aa1-41), the GAP domain (via interaction with active Rho), and the IG^232,233^ residues are important for cortical F-actin bundling and colocalization, whereas at least 450aa of the IDR at the carboxyl-terminus is needed for proper colocalization with cortical F-actin.

### RGA-3/4 directly binds to and bundles F-actin *in vit*ro

The above results demonstrated that RGA-3/4 colocalizes with and bundles F-actin *in vivo* and identified regions of the protein necessary for these activities. However, they did not determine whether RGA-3/4-F-actin interaction is direct or indirect. To distinguish between these possibilities, we expressed and purified recombinant full-length RGA-3/4-GFP protein (Figure 4B). Successful purification required the GFP tag and storage at low protein concentrations under high arginine and salt conditions to maintain solubility. To minimize RGA-3/4 aggregation during incubation with F-actin, we developed a bulk *in vitro* F-actin binding and bundling assay: in brief, purified RGA-3/4-GFP was incubated with *in vitro*-assembled F-actin, quickly mixed with low-melt agarose which was induced to gel by chilling (to immobilize the RGA-3/4-F-actin mixture), and then imaged within the gel using confocal microscopy (Figure 4A). This approach revealed that RGA-3/4 bound to F-actin and induced F-actin bundling similar to its behavior in *Xenopus* oocytes (Figure 4C). Because *in vitro* bundled F-actin concentrates filament signal within a smaller area, F-actin bundling was measured as the mean F-actin fluorescence intensity per unit area. Compared to the control, RGA-3/4-GFP significantly increased F-actin bundling (Figure 4D). To quantify F-actin binding, the number of pixels containing overlapping F-actin and RGA-3/4-GFP signal was measured; to distinguish between real and coincidental overlapping signal, the F-actin channel was rotated 90° to the left and colocalization was remeasured. Rotation significantly reduced the number of overlapping pixels, indicating that RGA-3/4 binding to F-actin was specific and not due to random spatial overlap (Figure 4E).

**Figure 4.**
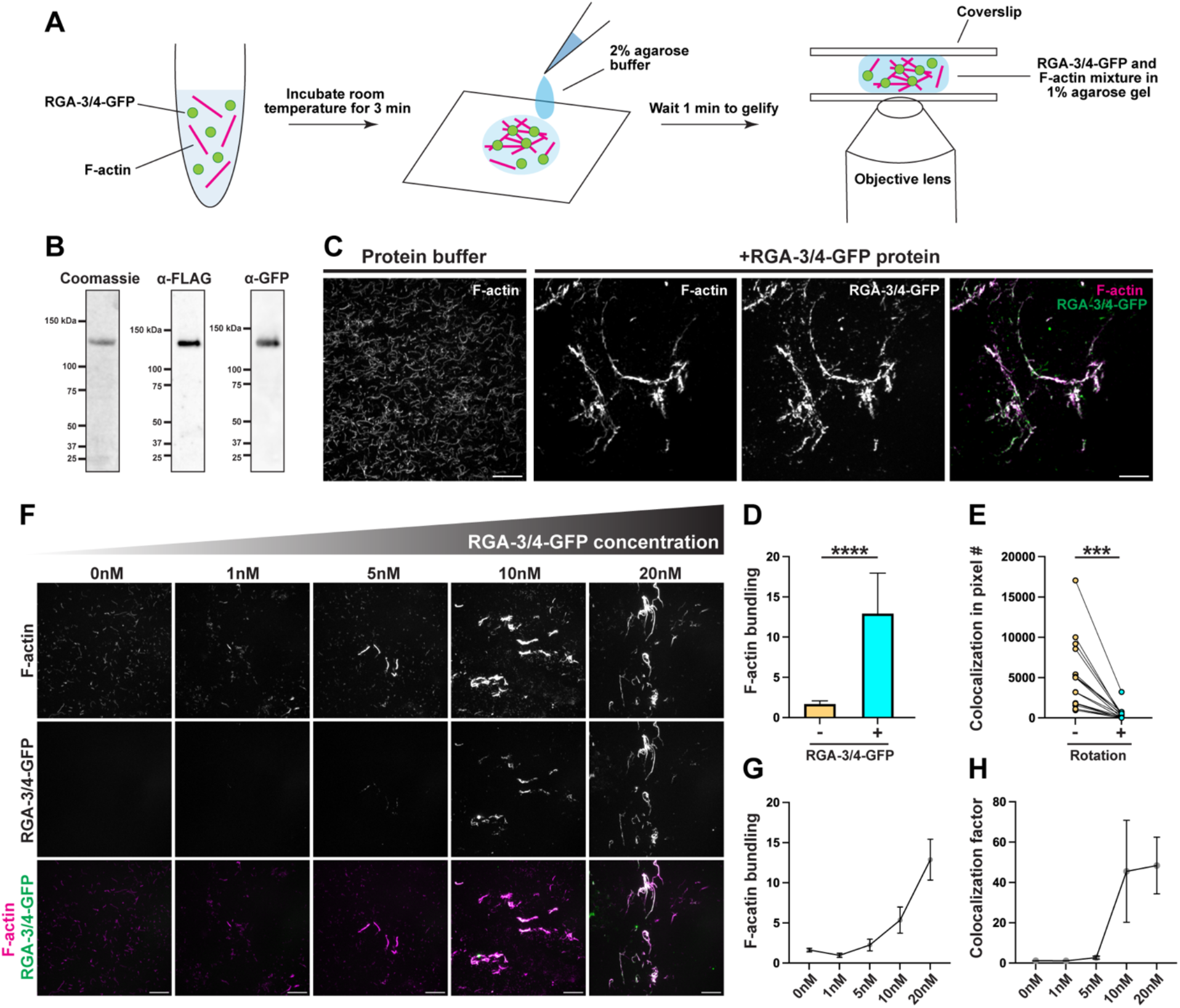
RGA-3/4 directly binds and bundles F-actin in vitro. (A) Schematic diagram of the experimental workflow. (B) Coomassie and immunoblot gels of purified Flag-Xenopus RGA-3/4-GFP protein. (C) F-actin incubated with protein buffer (control) or 20nM of purified RGA-3/4-GFP protein. F-actin was labelled with Alexa546-phalloidin and used at 30nM final concentration. Scale bar: 20 *μ* m. (D) Quantification of F-actin bundling. Measurements from 5 experiments. ****, <0.0001. Mean with 95%CI is shown. Unpaired t-test was performed for statistical analysis. (E) Quantification of colocalized pixel numbers without and with rotating F-actin image 90° to the left. Measurements from 5 experiments. ***=0.0002. Paired t-test was performed for statistical analysis. (F) F-actin incubated with increasing concentration of purified RGA-3/4-GFP protein. Scale bar: 20 *μ*m. Quantification of (G) F-actin bundling and (H) colocalization factor. Measurements from 5 experiments. Mean with SEM is shown.

To further characterize the interaction between RGA-3/4 and F-actin, we next asked how the interaction depends on RGA-3/4-GFP concentration. The results show that increasing the RGA-3/4-GFP concentration produced non-linear increases in both F-actin binding and bundling (Figure 4, F-H). Collectively, these results reveal that RGA-3/4 directly binds to and bundles F-actin.

### Structure-function analysis of RGA-3/4 interaction with F-actin *in vitro*

To test whether the regions required for cortical F-actin interaction in oocytes were also required for F-actin interaction *in vitro*, we performed *in vitro* structure-function analysis using purified RGA-3/4 mutants based on the *in vivo* analysis above. The aa42-801 mutant, which lacked detectable cortical F-actin bundling activity and colocalization *in vivo* (Figure 3, B-D), also failed to interact with F-actin *in vitro* (Figure 5, C-E). The I232A,G233A mutant also showed impaired F-actin interaction *in vivo* (Figure 3, B-D; Supplemental Figure S2). Consistent with this, I232A,G233A showed an attenuated F-actin interaction phenotype *in vitro* (Figure 5, C-E). The carboxyl-terminal truncations – aa1-350, aa1-300, aa1-250 – all induced cortical F-actin bundling but showed diminished colocalization *in vivo* (Figure 3, B-D). *In vitro*, these mutants also retained F-actin bundling activity and showed decreased binding to F-actin (Figure 5, C-D). Strikingly, the *in vitro* assay revealed a distinct localization pattern that may explain the diminished colocalization phenotype *in vivo*: while full-length RGA-3/4-GFP extensively and continuously decorated bundled F-actin, the carboxyl-terminal truncations localized to bundled F-actin in discrete puncta (Figure 5C). Taken together, the *in vitro* structure-function analysis parallels the *in vivo* analysis, indicating that the amino-terminal region and IG^232,233^ residues are important for F-actin interaction while half of the carboxyl-terminal region is needed for extensive RGA-3/4 localization along F-actin and is dispensable for bundling.

**Figure 5.**
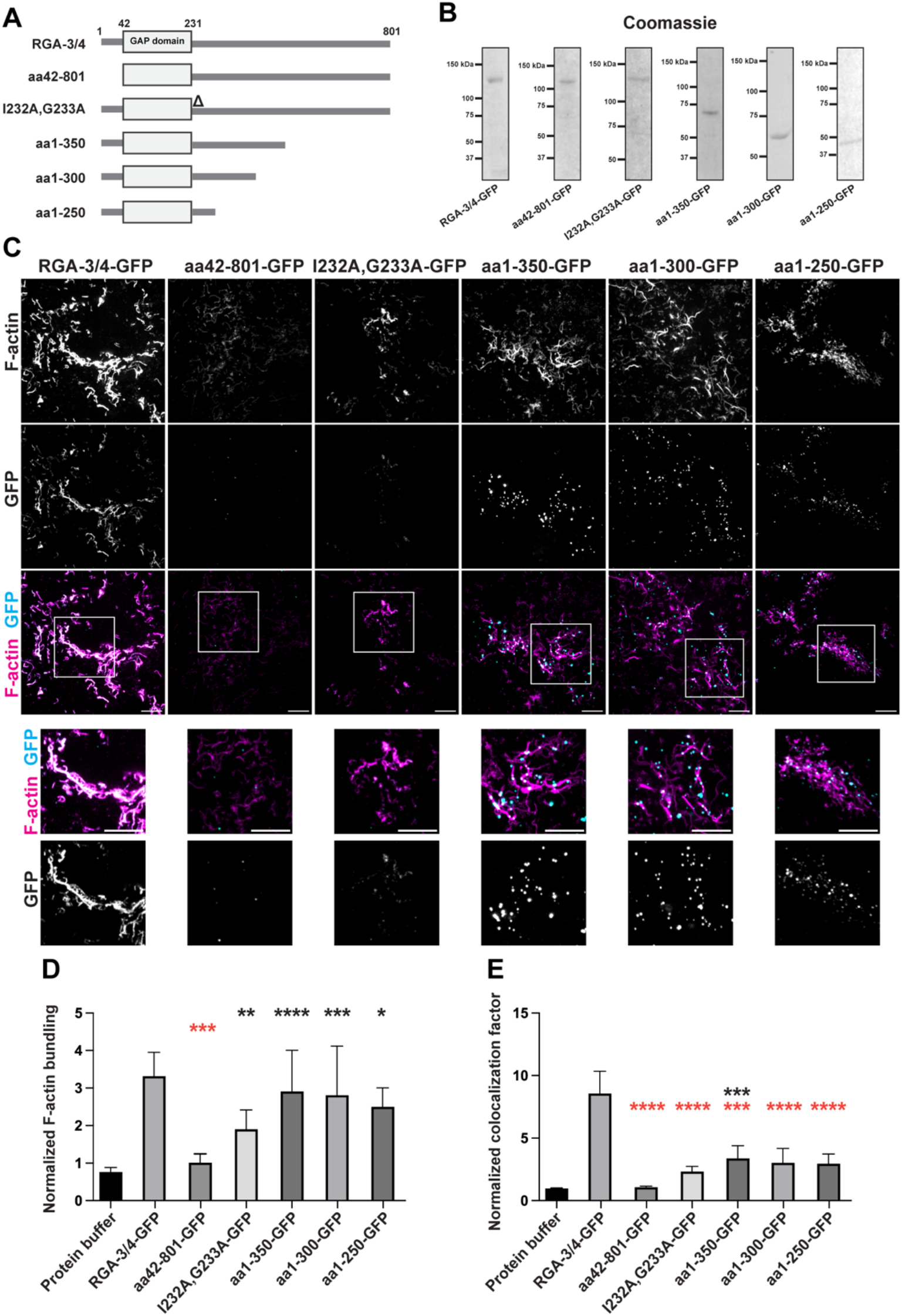
Structure-function analysis of RGA-3/4 interaction with F-actin in vitro. (A) Schematic diagrams of RGA-3/4 and various truncated or mutated constructs. (B) Coomassie gels of purified proteins of constructs in A. (C) Alexa546-phalloidin-labelled F-actin incubated with wildtype, truncated, or mutated RGA-3/4-GFP proteins. F-actin was used at 30nM final concentration and all RGA-3/4-GFP proteins were used at 20nM final concentration. Scale bar: 20 *μ* m. (D) Quantification of normalized F-actin bundling. *=0.0262; **=0.0053; ***=0.0003 (red asterisk); ***=0.0006 (black asterisk); ****, <0.0001. (E) Quantification of normalized colocalization factor. ***=0.0001 (red asterisk); ***=0.0003 (black asterisk); ****, <0.0001. (D and E) Measurements from 15 experiments. Mean with 95%CI is shown. One-way ANOVA with Dunnett post-hoc test for multiple comparisons was performed for statistical analysis. Black asterisks: comparison to protein buffer (negative control). Red asterisks: comparison to RGA-3/4-GFP (positive control).

### RGA-3/4 cooperatively binds to single actin filaments

The *in vitro* bulk F-actin assay demonstrated that RGA-3/4 directly binds to and bundles F-actin (Figure 4). However, this assay does not reveal how RGA-3/4 interacts with individual actin filaments. The non-linear concentration dependence of RGA-3/4-F-actin interaction suggested that RGA-3/4 enrichment on F-actin may be cooperative. A hallmark feature of cooperative F-actin binding is spatial heterogeneity, rather than uniform enrichment along the filament (Sharma *et al*., 2012; Hayakawa *et al*., 2014; Schmidt *et al*., 2015; Hosokawa *et al*., 2021). To determine whether RGA-3/4 binds individual filaments cooperatively, RGA-3/4-GFP recruitment to F-actin was visualized using time-lapse TIRF microscopy. Consistent with the results from the bulk *in vitro* assay, RGA-3/4-GFP specifically colocalized with actin at the single-filament level (Figure 6B). Time-course imaging showed that RGA-3/4-GFP progressively enriched on single actin filaments over time (Figure 6C) and did so in a discontinuous manner, as expected for cooperative binding. To examine the non-uniform spatial pattern of RGA-3/4-GFP localization, kymographs of individual filament were generated. This analysis confirmed that RGA-3/4-GFP signal increased over time in distinct regions within the filament, remaining spatially separated with neighboring regions (Figure 6E). Line-scan intensity profiles of RGA-3/4-GFP along the filament further revealed that instead of producing a single broad intensity peak along the filament, RGA-3/4-GFP formed multiple discrete intensity peaks that increased over time while maintaining their heterogeneous spatial pattern (Figure 6, F-F”).

**Figure 6.**
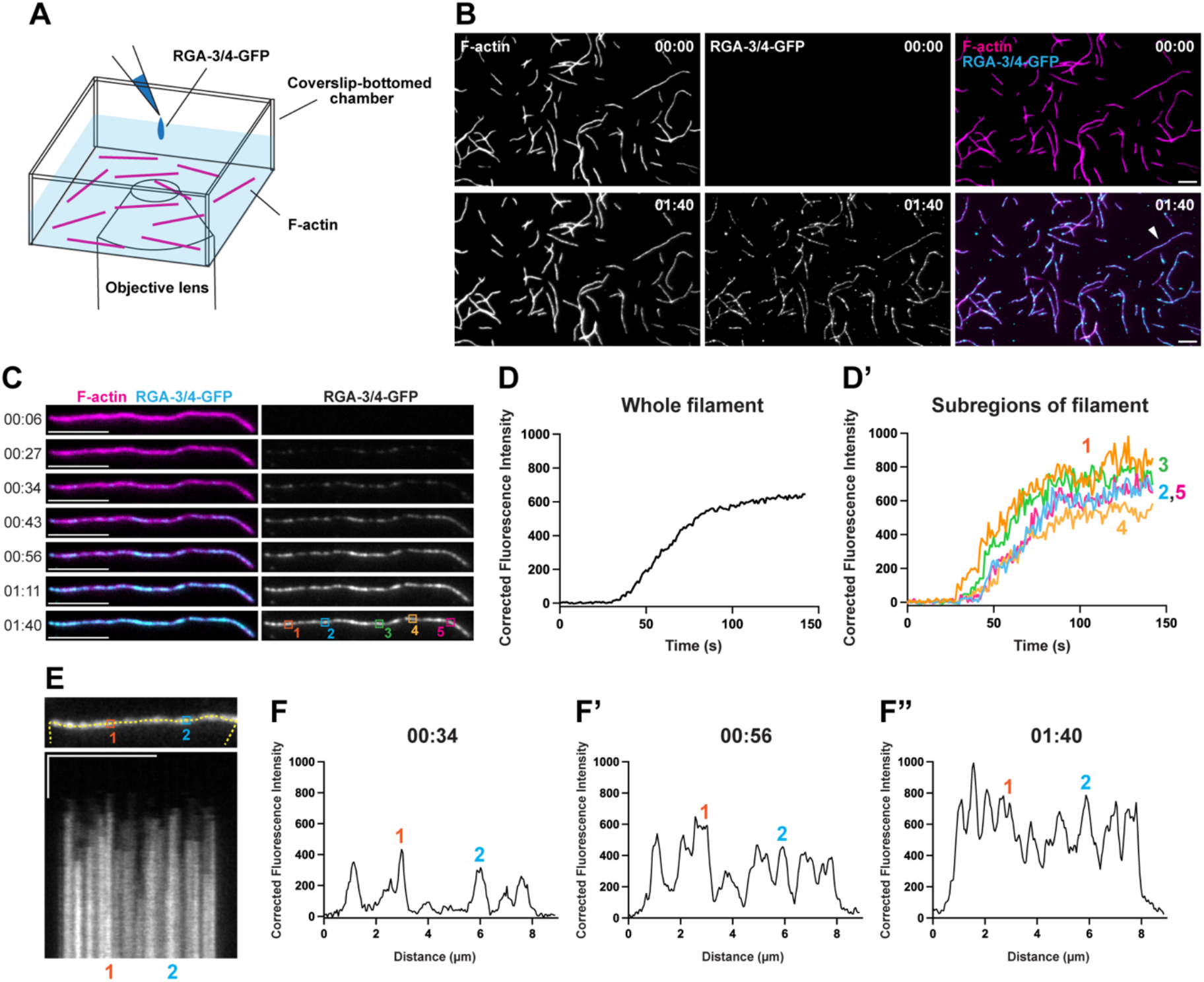
RGA-3/4 cooperatively binds to single actin filaments. (A) Schematic diagram of experimental setup. (B) Representative still images of F-actin before and after RGA-3/4-GFP enrichment. Arrowhead indicates F-actin shown in C. (C) Time-course montage of RGA-3/4-GFP accumulation on F-actin. Corrected fluorescence intensity over time of RGA-3/4-GFP measured along (D) whole filament and (D’) subregions of the filament. (E) Region of the filament shown in C, outlined with where kymograph was generated (top), and the corresponding kymograph (bottom). (F-F”) Line-scan intensity profile of RGA-3/4-GFP along the filament shown in E, at different time points. X scale bar: 5*μ*m. Y scale bar: 30 seconds.

Large, extremely bright RGA-3/4-GFP clusters occasionally appeared on the background or landed directly on F-actin within the TIRF field (Supplemental Figure S3); this behavior became more common at later time points in the experiment or at higher RGA-3/4-GFP concentration as expected if some of the protein was aggregating in the low-salt physiological buffer used for TIRF imaging (Supplemental Figure S3; see also Methods for protein purification). When clusters landed on F-actin, the intensity traces showed abrupt increases in fluorescence intensity (Supplemental Figure S3, A’, B’ and B”). Because these abrupt landing events do not reflect RGA-3/4-GFP progressive enrichment on F-actin, we excluded obvious aggregate-landing events from subsequent measurements.

To determine how the spatially heterogeneous pattern of RGA-3/4 developed over time, we measured RGA-3/4-GFP fluorescence intensity along the whole filament over time. The RGA-3/4-GFP fluorescence exhibited a sigmoidal accumulation profile (Figure 6D), supporting the interpretation of cooperative binding of RGA-3/4 to F-actin. We additionally measured individual subregions along the filament to more accurately capture local differences in accumulation pattern of RGA-3/4-GFP on F-actin over time. These subregion measurements still produced a sigmoidal enrichment profile (Figure 6D’), indicating that the cooperative F-actin binding is a characteristic feature of RGA-3/4.

### RGA-3/4 enrichment on F-actin precedes filament crosslinking

The standard TIRF set-up with the actin filaments anchored to the coverslip does not lend itself to observation of crosslinking or bundling events. However, we noticed that during live TIRF imaging, filaments that were incompletely anchored to the coverslip displayed rapid rotational or flexural movements. In the absence of RGA-3/4-GFP, these filaments continued through the time of imaging (data not shown). In contrast, in the presence of RGA-3/4, F-actin crosslinking and bundling of incompletely anchored filaments were observed when the mobile filaments were within reach of other filaments. To determine how these events relate to RGA-3/4 binding to F-actin, we examined the timing of RGA-3/4-GFP localization relative to filament crosslinking. This analysis showed that RGA-3/4-GFP localized to one or both of the actin filaments involved prior to crosslinking events (60/61 events). Two examples of this behavior are shown in Figure 7. In Figure 7A, a loosely attached actin filament rotates around with one of its ends apparently brushing a nearby, immobile filament until it forms a stable, end-on attachment at the site of RGA-3/4 accumulation. In Figure 7B, a filament anchored at only one end is positioned between two immobile filaments; this filament rotates freely past the upper filament until it eventually forms a stable, parallel attachment to the lower filament at points where RGA-3/4 has accumulated on the latter. These observations directly document the process of RGA-3/4-dependent crosslinking inferred from both the *in vivo* and the bulk *in vitro* assays.

**Figure 7.**
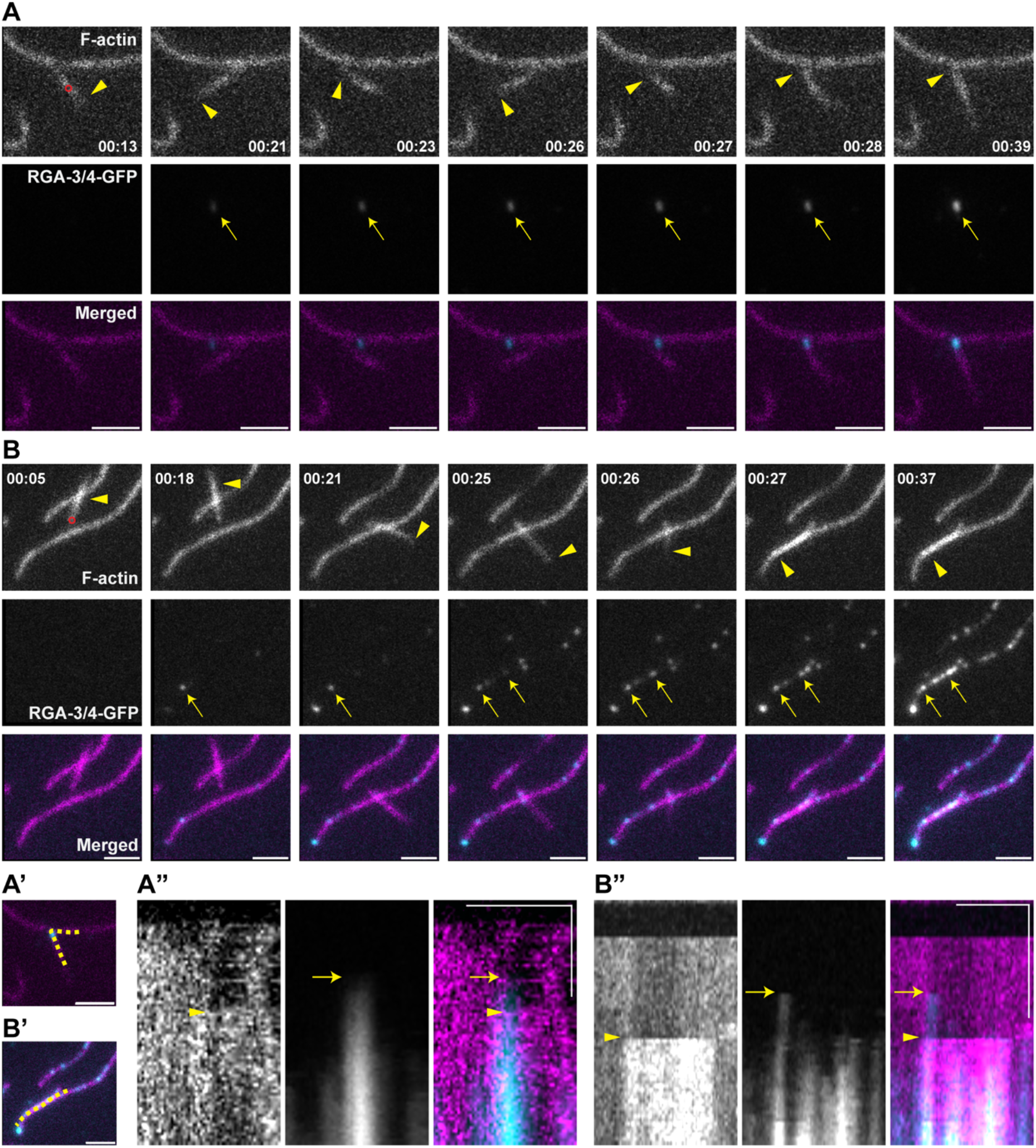
Direct visualization of filament crosslinking. Time-lapse images showing RGA-3/4-GFP enriching at the sites of F-actin (A) crosslinking or (B) bundling prior to the event. Red circle indicates point at which the filament is anchored. Merged images show F-actin in magenta and RGA-3/4-GFP in cyan. Arrowheads indicate F-actin that subsequently get crosslinked or bundled. Arrows indicate RGA-3/4-GFP localization. (A’ and B’) Images shown in A and B, outlined with where kymographs were generated. (A” and B”) Corresponding kymographs showing RGA-3/4-GFP and F-actin dynamics over time. X scale bar: 2*μ*m. Y scale bar: 30 seconds.

### Structure-function analysis of RGA-3/4 interaction with single actin filaments

RGA-3/4-GFP rapidly accumulated on individual actin filaments in spatially heterogeneous patterns that collectively decorated much of the filament. To determine which regions of RGA-3/4 are responsible for this behavior, previously characterized RGA-3/4 mutants were analyzed with live TIRF imaging. The aa42-801 mutant did not show any appreciable localization to F-actin (Figure 8, C-C”). The I232A,G233A mutant showed attenuated enrichment on F-actin but its fluorescence intensity still increased over time with a discontinuous and sigmoidal profile (Figure 8, D-D”). The aa1-350 mutant accumulated as discrete puncta that sparsely occupied the filament (Figure 8, E and E”). Over time, aa1-350 fluorescence intensity reflected puncta slowly increasing in intensity, not reaching the plateau during the time course imaged (Figure 8, E and E’). Similar accumulation patterns were observed for further carboxyl-terminal truncation mutants (Supplemental Figure S4).

**Figure 8.**
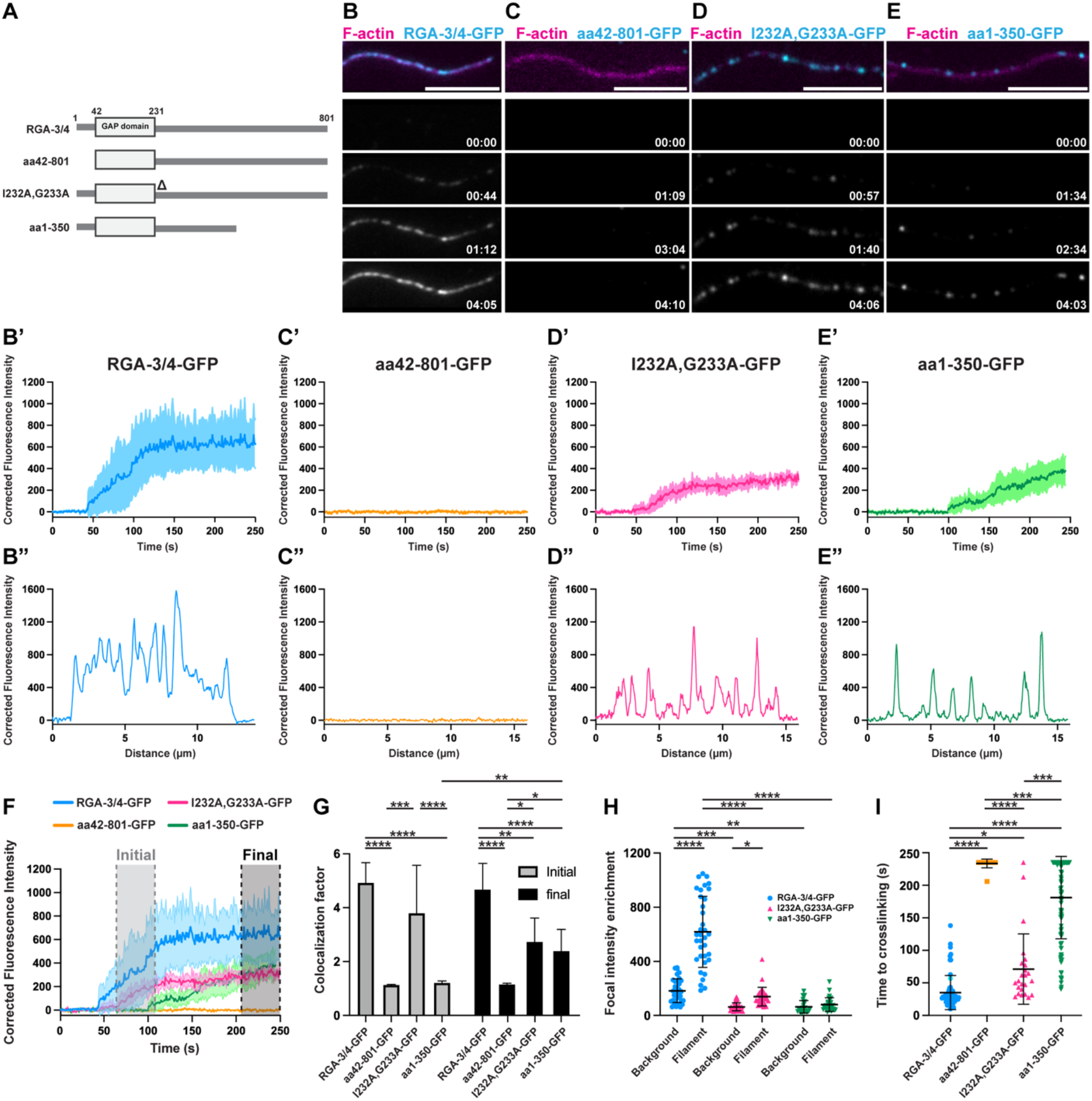
aa42-801, I232A,G233A and aa1-350 mutants show distinct defects in RGA-3/4 interaction with F-actin. (A) Schematic diagrams of RGA-3/4 and various truncated or mutated constructs. (B-E) Time-course montage of wildtype or mutant RGA-3/4-GFP proteins accumulating on individual filament. Scale bar: 5 *μ* m. (B’-E’) Corrected fluorescence intensity over time of wildtype or mutant RGA-3/4-GFP proteins measured along subregions of the filament. Mean±SD is shown. (B”-E”) Line-scan intensity profile of wildtype or mutant RGA-3/4-GFP proteins measured along the filament. (F) Overlay of graphs shown in B’-E’. (G) Quantification of colocalization between F-actin and wildtype or mutant RGA-3/4-GFP proteins at initial and final time points. Measurements from 8 experiments. *=0.0466 (aa42-801-GFP vs. I232A,G233A-GFP); *=0.0434 (aa42-801-GFP vs. aa1-350-GFP); **=0.0074 (RGA-3/4-GFP vs. I232A,G233A-GFP); **=0.0047 (aa1-350-GFP; initial vs. final); ***=0.0001; ****, <0.0001. Mean with 95%CI is shown. Two-way ANOVA with Tukey post-hoc test for multiple comparisons was performed for statistical analysis. (H) Quantification of wildtype or mutant RGA-3/4-GFP focal enrichment on background and filament. Measurements from 6 experiments. *=0.0262; **=0.0011; ***=0.0008; ****, <0.0001. (I) Quantification of the time required for F-actin crosslinking in the presence of wildtype or mutant RGA-3/4-GFP proteins. Measurements from 5 experiments. *=0.0164; ***=0.0003; ****, <0.0001. (H and I) Mean±SD is shown. Each dot represents measurement of a single filament. One-way ANOVA with Tukey post-hoc test for multiple comparisons was performed for statistical analysis.

To first analyze the timing and the extent of RGA-3/4 binding to F-actin, we measured the overlapping fluorescence signals of RGA-3/4-GFP and the mutants with single actin filaments at the initial and final time points (Figure 8; F and G). The aa42-801 mutant showed no detectable binding to F-actin at either the initial or final time points (Figure 8G). The I232A,G233A mutant exhibited F-actin binding at both time points, but the binding activity was significantly lower than that of RGA-3/4-GFP (Figure 8G). The aa1-350 mutant showed binding only at the final time point and this binding was attenuated relative to full-length RGA-3/4 (Figure 8G). Similar patterns were observed for further carboxyl-terminal truncation mutants (Supplemental Figure S4).

RGA-3/4-GFP rapidly accumulated at sites on F-actin after the initial binding compared to RGA-3/4-GFP that landed randomly on background regions. To analyze such behavior and determine whether F-actin preferentially promotes RGA-3/4 recruitment, we measured focal RGA-3/4-GFP intensity following initial binding on either F-actin or on the background. RGA-3/4-GFP showed significantly greater focal intensity enrichment on F-actin than on background (Figure 8H). I232A,G233A-GFP also showed significantly greater enrichment on F-actin compared to the background, but to a lesser extent than RGA-3/4-GFP (Figure 8H). In contrast, aa1-350-GFP showed no significant difference in enrichment between F-actin and background (Figure 8H). Interestingly, RGA-3/4-GFP also showed greater focal enrichment on background than either mutant, suggesting that RGA-3/4 may have an intrinsic capacity to self-recruit that is further enhanced upon binding to F-actin. Overall, these results suggest that distinct parts of RGA-3/4 regulate different aspects of its interaction with F-actin; the amino-terminal region is required for the initial F-actin binding, the IG^232,233^ residues promote full enrichment, and approximately half of the carboxyl-terminal region enables rapid and robust RGA-3/4 accumulation on F-actin.

RGA-3/4-GFP accumulation on F-actin was shortly followed by filament crosslinking. To test how mutations affecting F-actin accumulation alter crosslinking activity, we measured the time to filament crosslinking in the presence of wildtype and mutant RGA-3/4-GFP. Filaments that were not crosslinked during the 240-second imaging window were scored as requiring 240 seconds, thus the analysis underestimates the time required for crosslinking by the mutants. The aa42-801 mutant largely failed to induce filament crosslinking, with most filaments remaining mobile and not crosslinked by 240-second (Figure 8I). The I232,G233A mutant induced filament crosslinking but exhibited delayed timing relative to wildtype RGA-3/4-GFP (Figure 8I). The aa1-350 mutant displayed a more pronounced delay, with a subset of filaments failing to undergo crosslinking (Figure 8I).

## Discussion

Here, we examined RGA-3/4-F-actin interactions using complementary *in vivo* and *in vitro* approaches. In *Xenopus* oocytes, RGA-3/4 colocalized with and bundled cortical F-actin. *In vitro*, purified recombinant RGA-3/4 directly bound to and bundled F-actin, demonstrating that these activities are intrinsic to RGA-3/4. Time-lapse TIRF imaging revealed that RGA-3/4 accumulated discontinuously along individual actin filaments and displayed rapid, sigmoidal enrichment over time, behaviors characteristic of cooperative binding (Sharma *et al*., 2012; Hayakawa *et al*., 2014; Schmidt *et al*., 2015; Hosokawa *et al*., 2021). Notably, RGA-3/4 enrichment preceded filament crosslinking, showing that RGA-3/4 binding actively promotes subsequent F-actin reorganization. Together, these findings establish RGA-3/4 as a direct, cooperative F-actin binding and bundling protein and reveal a previously unrecognized mechanism by which RGA-3/4 can regulate the actin cytoskeleton independently of its ability to inactivate Rho.

Structure-function analysis identified the aa1-41 region, conserved IG^232,233^ residues, and approximately half of the carboxyl-terminal region as parts of RGA-3/4 contributing to different aspects of the F-actin interaction (Figure 9A). First, the aa1-41 region was sufficient for cortical F-actin colocalization, and deleting the region (aa42-801) abolished all detectable F-actin interactions both *in vivo* and *in vitro*. Although the aa1-41 region lacks recognizable F-actin binding motifs, AlphaFold structure prediction suggests that it contains an amphipathic alpha-helical structure. Well-characterized F-actin probes such as F-tractin and Lifeact have been shown to bind to F-actin via an amphipathic helix rather than using a canonical actin binding motif (Belyy *et al*., 2020; Shatskiy *et al*., 2025). The aa1-41 region of RGA-3/4 may therefore act as a direct F-actin binding site, in which hydrophobic residues positioned along one side of an amphipathic helix contact the hydrophobic pocket of actin subunits.

**Figure 9.**
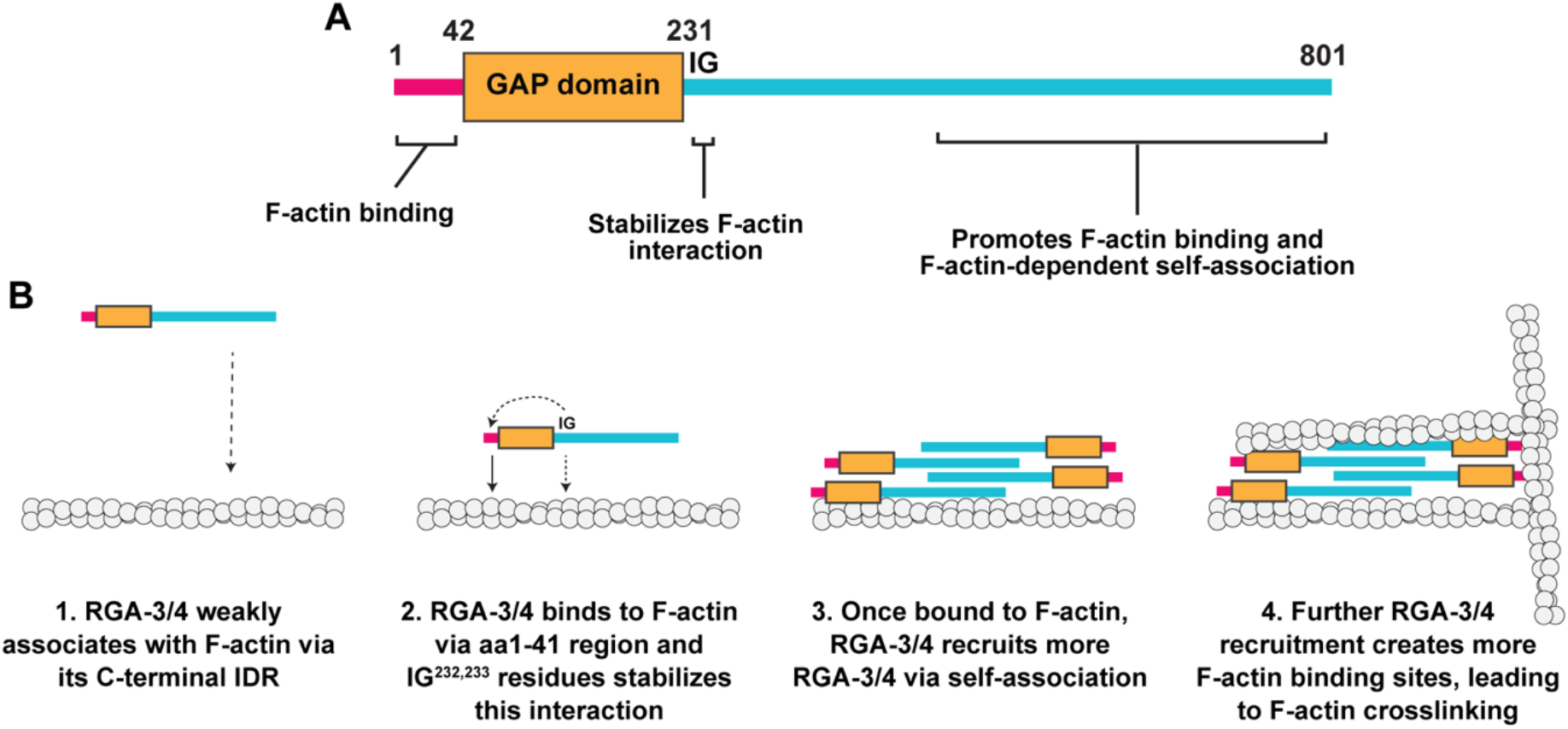
Schematic summary of RGA-3/4 interaction with F-actin. (A) Schematic summary of RGA-3/4 regions and their proposed contributions to RGA-3/4-F-actin interaction. (B) Hypothesized model for stepwise RGA-3/4 binding and bundling F-actin *in vitro*.

The structure-function analysis yielded the surprising result that while the I232A,G233A fails to show detectable F-actin interaction in *Xenopus* oocytes, it nonetheless displays diminished yet significant F-actin binding and bundling activity *in vitro*. In principle, this discrepancy could be explained if this mutation impairs RGA-3/4 folding or reduces the stability of the RGA-3/4-F-actin interaction. Although these possibilities cannot be excluded, the fact that it binds and bundles F-actin better than the aa1-350 mutant – which can bundle cortical F-actin *in vivo* – suggests that the *in vivo* phenotype likely reflects additional cellular regulation. One possibility is that RGA-3/4 has a binding partner that modulates actin binding *in vivo* and the I232A,G233A mutation somehow alters the interaction with that partner. The obvious candidate partner is Rho, raising the possibility that the Rho GTP hydrolysis cycle is entrained to a cycle of F-actin binding and release *in vivo*, such that interaction with active Rho promotes F-actin binding while the resultant hydrolysis of GTP by Rho promotes release of F-actin by RGA-3/4. Such entrainment would explain the findings obtained with the two Rho GAP mutants: the R80A mutant, which is predicted to bind to Rho-GTP but be unable to promote GTP hydrolysis, actually shows significantly better colocalization with cortical F-actin than wild-type RGA-3/4, while the R80E mutant, which is predicted to be unable to bind to Rho-GTP, fails to colocalize with cortical F-actin. A role for the IG^232,233^ residues in regulating a cycle of RGA-3/4, Rho, and F-actin binding is also consistent with the position of these residues immediately adjacent to the GAP domain.

The structure-function analysis showed that the carboxyl-terminal truncation mutants – aa1-350, aa1-300, and aa1-250 – induced cortical F-actin bundling but showed reduced colocalization with cortical F-actin in the oocytes. This result was also initially puzzling: how can reduced F-actin interaction fail to reduce bundling? The *in vitro* findings provided a potential explanation. That is, *in vitro*, the aa1-350 mutant retained F-actin binding and bundling but exhibited a striking alteration in the pattern of F-actin binding relative to full-length RGA-3/4, such that the aa1-350 accumulated on filaments as discrete puncta after a prolonged delay. Further, unlike full-length RGA-3/4, aa1-350 failed to accumulate between the puncta and the puncta themselves displayed little additional accumulation after their initial formation. Thus, the loss of measurable colocalization of aa1-350 with cortical F-actin in spite of its ability to bundle cortical F-actin likely reflects the fact that the diminished puncta are still capable of crosslinking F-actin over the several hours of expression in the oocytes in spite of the fact that there is substantially less on the actin filaments. A similar argument applies to the aa1-300 and aa1-250 mutants. Thus, most of the carboxyl-terminal IDR is not strictly required for F-actin binding and bundling, but instead modulates the kinetics, localization pattern, and cooperative enrichment of RGA-3/4 on F-actin.

These findings are of particular interest because IDRs have emerged as important regulatory features that can support multivalent interactions, self-association, and phase separation (Brangwynne *et al*., 2015; Martin and Holehouse, 2020; Holehouse and Kragelund, 2024). Depending on the molecular and cellular context, these properties can produce diverse effects on protein localization, assembly, and activity (Holehouse and Kragelund, 2024). Thus, the carboxyl-IDR of RGA-3/4 could contribute to F-actin interaction through the intrinsic properties of IDR rather than, or in addition to a site that is specifically responsible for F-actin binding. For example, the cooperative binding of RGA-3/4 to F-actin reported here requires the IDR, which could imply that the IDR normally promotes oligomerization of RGA-3/4 following its initial binding to F-actin. Consistent with this possibility, IDRs in many proteins have been shown to promote intermolecular interactions and higher order self-assembly (Martin and Holehouse, 2020; Wiegand and Hyman, 2020). Moreover, the RGA-3/4 IDR was additionally required for rapid and discontinuous, yet complete, labeling along the filaments. Charged IDRs can promote electrostatic steering, accelerating association with oppositely charged binding partners without themselves forming the primary binding interface (Vuzman *et al*., 2010; Ganguly *et al*., 2013; Kristensen *et al*., 2018) and may subsequently facilitate diffusion along the binding surface (Bigman and Levy, 2023). Since the RGA-3/4 IDR is highly positively charged and F-actin presents a strongly negative surface, we propose that electrostatic steering increases the local concentration of RGA-3/4 near F-actin, facilitating rapid association and subsequent spreading along the filament. Thus, the carboxyl-IDR may promote RGA-3/4-F-actin interaction through complementary mechanisms: facilitating local enrichment through IDR-dependent self-association and enhancing F-actin association and spreading through electrostatic steering.

Although cortical F-actin bundling activity and colocalization were conserved between *Xenopus* and human RGA-3/4, human RGA-3/4 exhibited significantly greater colocalization with F-actin than *Xenopus* RGA-3/4 (Figure 1E). Human RGA-3/4 contains approximately 200 additional residues in its carboxyl-terminus (Figure 1A), extending the IDR and potentially strengthening IDR-mediated interactions with F-actin. In addition, a recent study identified a short linear motif within the human RGA-3/4 IDR that mediates low-affinity F-actin binding in other contexts (Paraschiakos *et al*., 2026); this motif is absent from the *Xenopus* RGA-3/4 IDR. These differences can further explain the structure-function analysis of human RGA-3/4, where the aa47-1023 and I238A,G239A showed impaired cortical F-actin bundling but still exhibited appreciable F-actin colocalization while the corresponding *Xenopus* mutants did not display any detectable F-actin interaction. Thus, the extended human RGA-3/4 IDR may provide additional F-actin interaction capacity that compensates for disruption of other F-actin binding sites.

Based on the combined results from this study, we propose a model for how RGA-3/4 binds to and accumulates on F-actin (Figure 9B). In this model, the carboxyl-terminal IDR promotes initial weak F-actin association, bringing RGA-3/4 in proximal distance to F-actin. RGA-3/4 then directly binds to F-actin through aa1-41 region, and IG^232,233^ residues stabilize this interaction. The carboxyl-terminal IDR subsequently promotes additional RGA-3/4 accumulation, converting the initial binding into cooperative enrichment and spreading on F-actin.

One of the major implications of this study is its potential to expand our current understanding of cortical excitability. Existing models assume that RGA-3/4 is recruited to F-actin in a simple linear fashion, where it functions primarily to inactivate Rho and provide negative feedback (Bement *et al*., 2015; Michaud *et al*., 2022). However, the findings presented here clearly indicate that the recruitment is more likely to be non-linear, both at the level of single filaments and as a consequence of F-actin bundling by RGA-3/4. To put it another way, the current findings suggest that RGA-3/4 and F-actin participate in a rapid positive feedback relationship. Incorporating these features of RGA-3/4 into current models of cortical excitability can therefore provide a more complete framework in understanding how negative feedback of cortical excitability is mediated. More broadly, because F-actin-dependent recruitment of Rho GAPs represent a recurring feature of negative feedback in multiple excitable systems (Bement *et al*., 2024), this study further motivates investigation into whether reciprocal feedback interactions between Rho GAP and F-actin are embedded within other cortical circuits and how such interactions contribute to system-specific cortical behaviors.

Since cortical excitability during cytokinesis directs the assembly of the actomyosin contractile ring that constricts the cell at the equatorial cortex, it raises the additional question of whether F-actin binding and bundling activity of RGA-3/4 contributes to F-actin organization within the contractile ring. In this study, RGA-3/4 enrichment preceded at the sites of crosslinking, and its subsequent interaction with F-actin was rapid and exponential. Such behavior suggests that a small localization of RGA-3/4 at the cell equator can provide enough sensitivity and cooperativity to strongly induce F-actin bundling within the short time window of cytokinesis. Future studies examining the F-actin binding and bundling activity of RGA-3/4 during cytokinesis may reveal an unconventional mechanism for how F-actin is organized within the contractile ring.

## Materials and Methods

### Constructs and mRNA

Plasmids for *Xenopus* RGA-3/4-GFP and various truncation plasmids were made from *Xenopus* RGA-3/4 (Michaud et al., 2022) by cloning the target region into pCS2+ plasmid with C-terminal GFP. Human RGA-3/4 clones (ArhGAP11a) were generated by amplifying cDNA (Horizon Discovery) via PCR and inserting the amplicon into appropriate vector plasmids (empty pCS2 or pCS2+ with C-terminal GFP). *Xenopus* RGA-3/4^I232A,G233A^-GFP was generated via PCR mutagenesis. Plasmids were linearized downstream of open reading frame via NotI-HF restriction enzyme digestion and were transcribed into mRNA via mMessage mMachine sp6 kit (Ambion, AM1340). All mRNAs were injected into *Xenopus* oocytes at needle concentration of 5.8*μ*M.

### Xenopus oocytes

About a quarter of the ovaries were extracted from adult female *Xenopus* laevis and rinsed in 1x Barth’s solution (87.4mM NaCl, 1mM KCl, 2.4mM NaHCO3, 0.82mM MgSO4, 0.6mM NaNO3, 0.7mM CaCl2, 10mM HEPES at pH7.4). The ovary chunks were finely cut and rinsed thoroughly with 1x Barth’s solution, followed by 8mg/mL collagenase treatment for 1hr at 16°C. Oocytes were then rinsed extensively with 1x Barth’s solution and allowed to recover at 16°C for at least 3hrs. After recovery, follicle cells were manually removed from stage VI oocytes, and the oocytes were injected with 40nl of mRNA. Oocytes were incubated at room temperature for 4-6hrs before fixation.

### Fixation

Oocytes were fixed in superfix solution (100mM KCl, 3mM MgCl, 10mM HEPES at pH 7.4, 150mM sucrose, 3.7% paraformaldehyde, 10% DMSO, 500mM ethylene glycol-bis(b-aminoethylether)-*N,N,N’,N’*-tetraacetic acid (EGTA), 0.2% Triton, and 2U/mL Alexa 546-Phalloidin) at room temperature for 30min. Following fixation, the oocytes were rinsed in 1x phosphate-buffered saline (PBS) for 30min and imaged shortly after.

### Protein purification

Proteins for *Xenopus* RGA-3/4-GFP and other mutants (aa42-801-GFP, I232A,G233A-GFP, aa1-350-GFP, aa1-300-GFP, aa1-250-GFP) were made via baculovirus system as previously described (Bement *et al*., 2015; Michaud *et al*., 2022). A 5’ terminal Kozak consensus sequence and FLAG epitope were added to the region of interest by PCR, and the resultant amplicon was inserted into pFastBac1 plasmid. The plasmid was transformed into DH10Bac EMBacY bacteria (Geneva Biotech MultiBac™) to generate recombinant bacmid DNA. Sf21 insect cells were transfected with the recombinant bacmid DNA, and infected with subsequent recombinant baculovirus to produce recombinant protein. For purification, insect cells were lysed in arginine-based solubilization buffer (500mM Arginine, 10mM MgCl2, 0.01mg/mL Dnase I, 10 *μ*g/mL Rnase, 1% Triton, 10 *μ*g/mL leupeptin, 2 *μ*g/mL aprotinin, 1*μ*g/mL pepstatin A, 40*μ*g/mL Phenylmethylsulfonyl fluorise (PMSF), 200*μ*g/mL benzamidine, 0.5*μ*g/mL E64, 5*μ*g/mL MG132, in 1x phosphate-buffered saline (PBS) pH 7.5) and the cell lysate was applied to anti-FLAG M2 affinity resin column (Sigma-Aldrich; #A2220-25mL). The protein was eluted with a low-pH arginine-based buffer (1M Arginine, 500mM KCl pH 4.4) and stored in high arginine and salt buffer (500mM Arginine, 500mM KCl, 25mM HEPES pH 7.5). Purification without a GFP tag or arginine-based solubilization buffer, as well as storage at high concentrations or without high arginine and salt concentrations, resulted in low protein yield, degradation, and aggregation.

### Immunoblot

Proteins were separated by SDS-PAGE (3-12% gradient gel) and transferred to nitrocellulose membrane. The membrane was blocked in blotto [5% (w/v) milk in 1x phosphate-buffered saline (PBS)] at room temperature for 1hr, followed by primary antibody incubation overnight at 4°C. Anti-FLAG M2 monoclonal antibody (Sigma, F1804) was used at 1:1000 dilution, and anti-GFP polyclonal antibody (Sigma, G1546) was used at 1:3000 dilution. The membrane was washed twice with excess 5% blotto at room temperature for 10min and incubated with secondary antibody at room temperature for 1hr. Anti-mouse donkey antibody (LI-COR, 926-32212) and anti-rabbit goat antibody (Invitrogen, A21076) were both used at 1:3000 dilution. The membrane was washed twice with 1x PBS for 10 min and was analyzed using Odyssey Fc Imaging System (LI-COR Biosystems).

### Bulk F-actin binding and bundling assay

F-actin bulk assay refers to the experimental workflow shown in Figure 4A. Actin from Cytoskeleton (AKL99) was used to polymerize F-actin as per the company’s guideline (Cytoskeleton Inc). 120U/mL Alexa 546-phalloidin-labelled F-actin was mixed with RGA-3/4-GFP proteins and incubated for 3min at room temperature. The mixture was then mounted on 2% BSA coated coverslip (22×22mm, no. 1.5, Globe Scientific Inc., 1404-15) and glass slide (Fisher, 125493). Final buffer concentration of the mixture was the following: 25mM HEPES pH 7.5, 5mM Tris-HCl, 0.2mM CaCl2, 2mM MgCl2, 140mM KCl, 1mM ATP, 125mM Arginine, 1% agarose. Final concentration of actin was 30nM and RGA-3/4-GFP proteins was 20nM unless noted otherwise in the figure legends.

### TIRF single actin filament assay

TIRF F-actin assay refers to the experimental setup shown in Figure 6A. Glass coverslips (22×60mm, no. 1.5, Alkali Scientific Inc., SM2108) were cleaned and functionalized as previously described (Bieling *et al*., 2010). Briefly, the coverslips were cleaned with 3M NaOH for 30min and subsequently with Piranha solution (3:2 ratio of sulfuric acid and 30% hydrogen peroxide) for 45min. The coverslips were then silanized with 3-(Glycidoxypropyl)trimethoxysilane (GOPTS) for 30min at 60 ° C. Coverslips were separated in acetone and functionalized with amine-PEG-biotin/amine-PEG-hydroxy (5:95 ratio in acetone) for 4hrs at 60°C. For chamber assembly, a functionalized coverslip was attached to sticky 18-well slide (Ibidi, 81818). Each well was blocked for 5min with blocking buffer (10mM imidazole, 100mM KCl, 1.5mM MgCl2, 1mM EGTA, 0.1mg/mL *κ*-casein, 1mM TCEP, 1% Pluronic F-127 pH 7.5), followed by 3min incubation with 75nM streptavidin and another 3min incubation with 600U/mL Alexa647-phalloidin-labelled biotinylated F-actin. 2 washes were done in between each step with 1x actin polymerizing buffer (1xAPB; 0.2mM CaCl2, 2mM MgCl2, 50mM KCl, 0.5mM DTT, 1mM ATP, 30mM HEPES pH 7.5). 1-3uL of RGA-3/4-GFP protein was added to each well for 100uL final volume of 1xAPB for live imaging.

### Microscopy and image processing

Fixed oocytes were imaged using a Prairie View Laser Scanning Confocal on a Nikon Eclipse Ti base (Bruker Nano surfaces) using 60x 1.4-NA oil objective. Oocytes were mounted on custom metal slides between coverslips (22×22mm, no. 1.5, Globe Scientific Inc., 1404-15) in 1x PBS. For F-actin bulk assay, a Prairie View Swept Field Confocal on a Nikon Eclipse Ti base (Bruker Nano Surfaces) was used with 60x 1.4-NA oil objective. TIRF imaging was performed using a Nikon Eclipse Ti2 base with iLas TIRF module using 100x 1.45-NA Plan Apo TIRF objective lens. Z-series were acquired using 0.5*μ*m-step in confocal while single z-plane was acquired for TIRF images.

All imaging processing was conducted using ImageJ/FIJI. Topdown confocal images in Figure 1B, 2B, 3B, 4C, 4F, 5C, S1A, and S2B are shown in maximum intensity projections. For sideview profiles in Figure 1D, 2B’, 3B’, and S2B’, images were processed using 3D projection function with brightest point projection method. Still frames from TIRF imaging shown in Figure 6B, 6C, 7A, 7B, 8B-E, S3A-C, and S4B were from movies registered for 2D drift using StackReg plugin and corrected for bleaching by using an exponential fit. Kymograph in Figure 6E and 7A”-B” was generated by reslicing the time-lapse images along a 1-px-thick line drawn across the field of view at location indicated in the figure. Kymograph was scaled in the y-axis using bicubic interpolation for display in the figures.

### Image quantifications

All image measurements were conducted using ImageJ/FIJI. Cortical F-actin bundling shown in Figure 1C, 2C, 3C, S1B, and S2C was calculated in reference to previous study (Sun *et al*., 2024). Mean signal intensity and standard deviation of cortical F-actin was measured, and F-actin bundling was calculated by dividing standard deviation with mean. For colocalization measurement in oocytes in Figure 1E, 2D, 3D, and S2D, F-actin and GFP signals were thresholded in 3D projected images, and colocalization ratio was calculated by dividing colocalized F-actin and GFP signals with the total GFP signals. Colocalized F-actin and GFP signals were measured using color counter plugin. Cortical F-actin bundling and colocalization ratio were normalized by the average of control (GFP alone) in each experiment.

*In vitro* F-actin bundling shown in Figure 4D, 4G, and 5D was quantified by first outlining F-actin using a mask created by thresholding F-actin signals, and dividing the mean signal intensity by the unit area of the mask. Colocalized pixel numbers of F-actin and GFP signals (from color counter plugin) were measured without and with rotating F-actin image 90° to the left. *In vitro* colocalization factor shown in Figure 4H, 5E, 8G, and S4D-D”, S4E was calculated by dividing the colocalized signals without rotation by the signals with rotation to eliminate any random colocalized signals (Figure 4E).

Fluorescence intensity over time was measured in 1-px-thick line drawn along the filament for whole filament in Figure 6D and S3B’, and in 2-px square box drawn on subregions of the filament in Figure 6D’, 8B’-E’, 8F, S3A’, S3B”, and S4C-C” using plot z-axis function. Plot profile function was used to measure fluorescence intensity across distance as shown in Figure 6F-F” and 8B”-E”. For each measurement, the fluorescence intensity value was corrected by subtracting the fluorescence intensity of regions outside the filament at the corresponding time point or distance.

To quantify intensity enrichment on F-actin in Figure 8H, fluorescence intensity over time was measured in 2-px square box positioned at sites where wildtype or mutant RGA-3/4-GFP accumulated on F-actin or background regions. For each measurement, baseline intensity was defined as the mean fluorescence before the onset of RGA-3/4-GFP accumulation, and plateau intensity as the mean fluorescence after enrichment reached a stable maximum. When enrichment did not reach a stable maximum, the fluorescence intensity at the final time point of the movie was used as the plateau intensity. Intensity enrichment was calculated by subtracting the baseline intensity from the plateau intensity. The intensity enrichment on F-actin was calculated by dividing the intensity enrichment on each filament by the mean intensity enrichment on the background regions within the corresponding movie.

For measurement of time to crosslinking in Figure 8I, TIRF movies were first analyzed to identify filaments that were only partially anchored on the coverslip and were within reach of other filaments. Filaments were considered crosslinking if they subsequently contacted another filament, ceased movement at the site of contact, and remained immobilized at the site of contact for at least 10 successive 1s frames. Crosslinking analysis was limited to the first 4 min of movies (due to photobleaching which made it difficult to unambiguously characterize filament movement), thus, any filament that remained mobile after 4 min was counted as lasting 240s.

## Acknowledgments

We thank Kurt Weiss and the University of Wisconsin-Madison Biochemistry Optical Core for the assistance in using TIRF microscopy. We also thank Peter Bieling for conceptual input and guidance in TIRF imaging experiments. This work was supported by awards from the National Science Foundation (MCB-2132606; MCB-2526692).

---

**Supplemental Figure S1.**
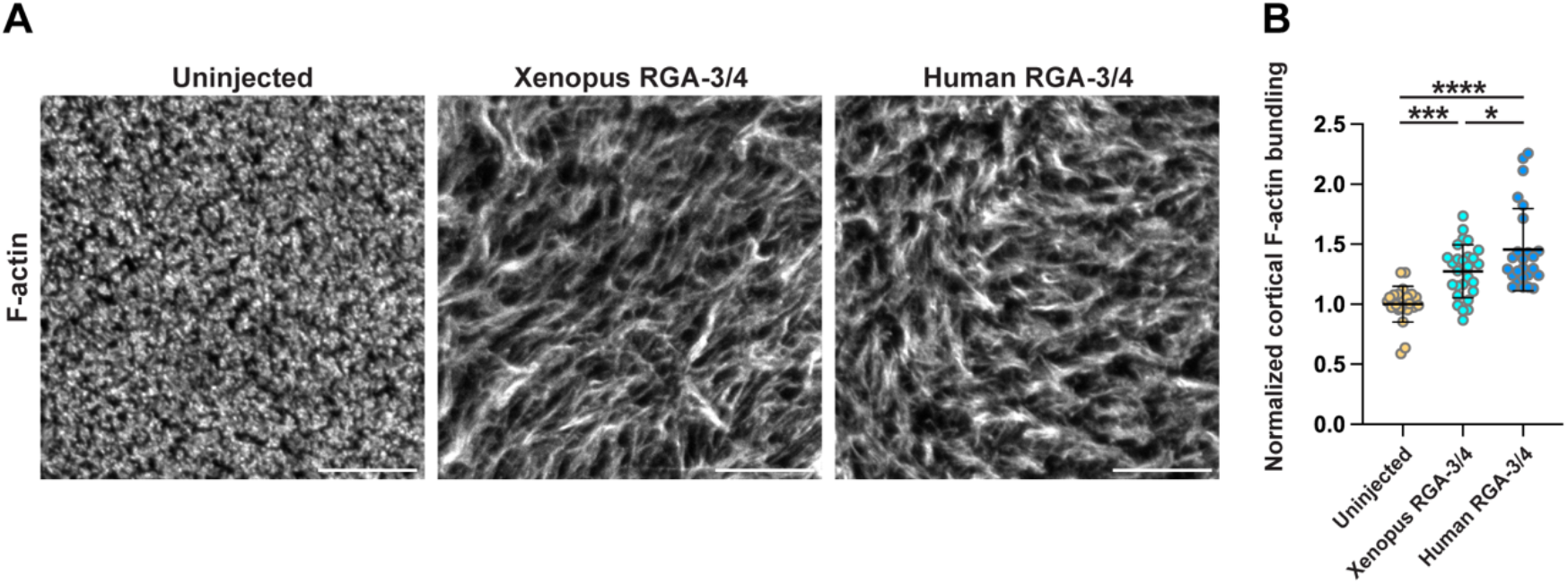
Untagged Xenopus and Human RGA-3/4 bundle cortical F-actin in immature Xenopus oocytes. (A) Cortex of uninjected immature Xenopus oocytes (control) and oocytes expressing Xenopus RGA-3/4 or Human RGA-3/4. The oocytes were fixed with Alexa546-phalloidin. Scale bar: 20*μ*m. (B) Quantification of normalized cortical F-actin bundling. Uninjected, n=24; Xenopus RGA-3/4, n=27; Human RGA-3/4, n=25; 3 experiments. *=0.0317; ***=0.0006; ****, <0.0001. One-way ANOVA with Tukey post-hoc test for multiple comparisons was performed for statistical analysis.

**Supplemental Figure S2.**
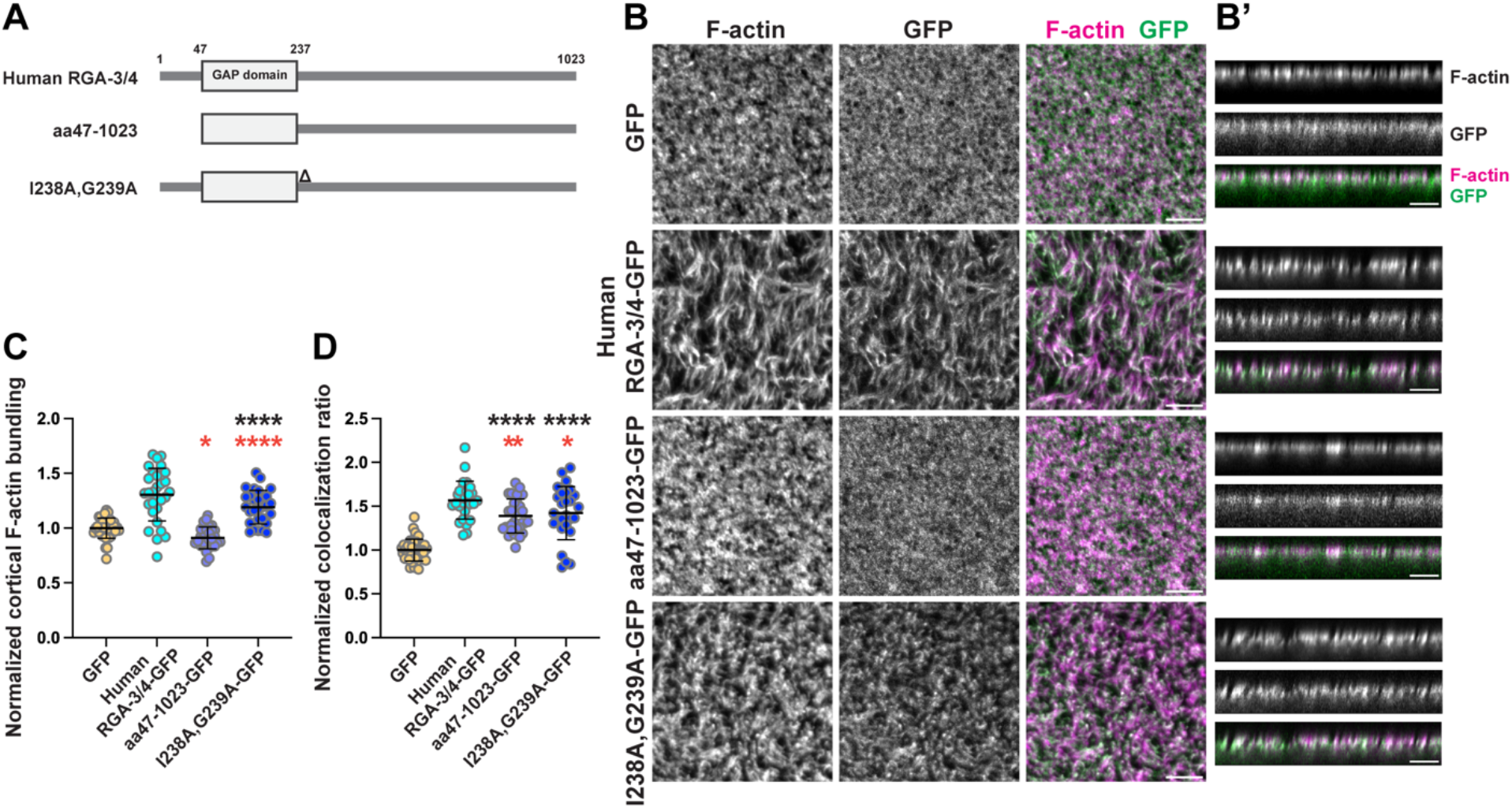
Structure-function analysis of Human RGA-3/4 interaction with cortical F-actin. (A) Schematic diagrams of Human RGA-3/4 and truncated or mutated constructs. (B) Representative oocyte from each treatment group. En-face and (B’) sideview profiles of the cortex are shown. The oocytes were fixed with Alexa546-Phalloidin. Scale bar: 10*μ*m. (C) Quantification of normalized cortical F-actin bundling. GFP, n=38; Human RGA-3/4-GFP, n=30; aa47-1023-GFP, n=29; I238A,G239A-GFP, n=28; 4 experiments. *=0.0152. ****, <0.0001. (D) Quantification of normalized colocalization ratio. Control, n=38; Human RGA-3/4-GFP, n=28; aa47-1023-GFP, n=32; I238A,G239A-GFP, n=27; 4 experiments. *=0.0280; **=0.0036. ****, <0.0001. (C and D) Mean±SD is shown. Each dot represents measurement in a single oocyte. One-way ANOVA with Dunnett post-hoc test for multiple comparisons was performed for statistical analysis. Black asterisks: comparison to GFP (negative control). Red asterisks: comparison to RGA-3/4-GFP (positive control).

**Supplemental Figure S3.**
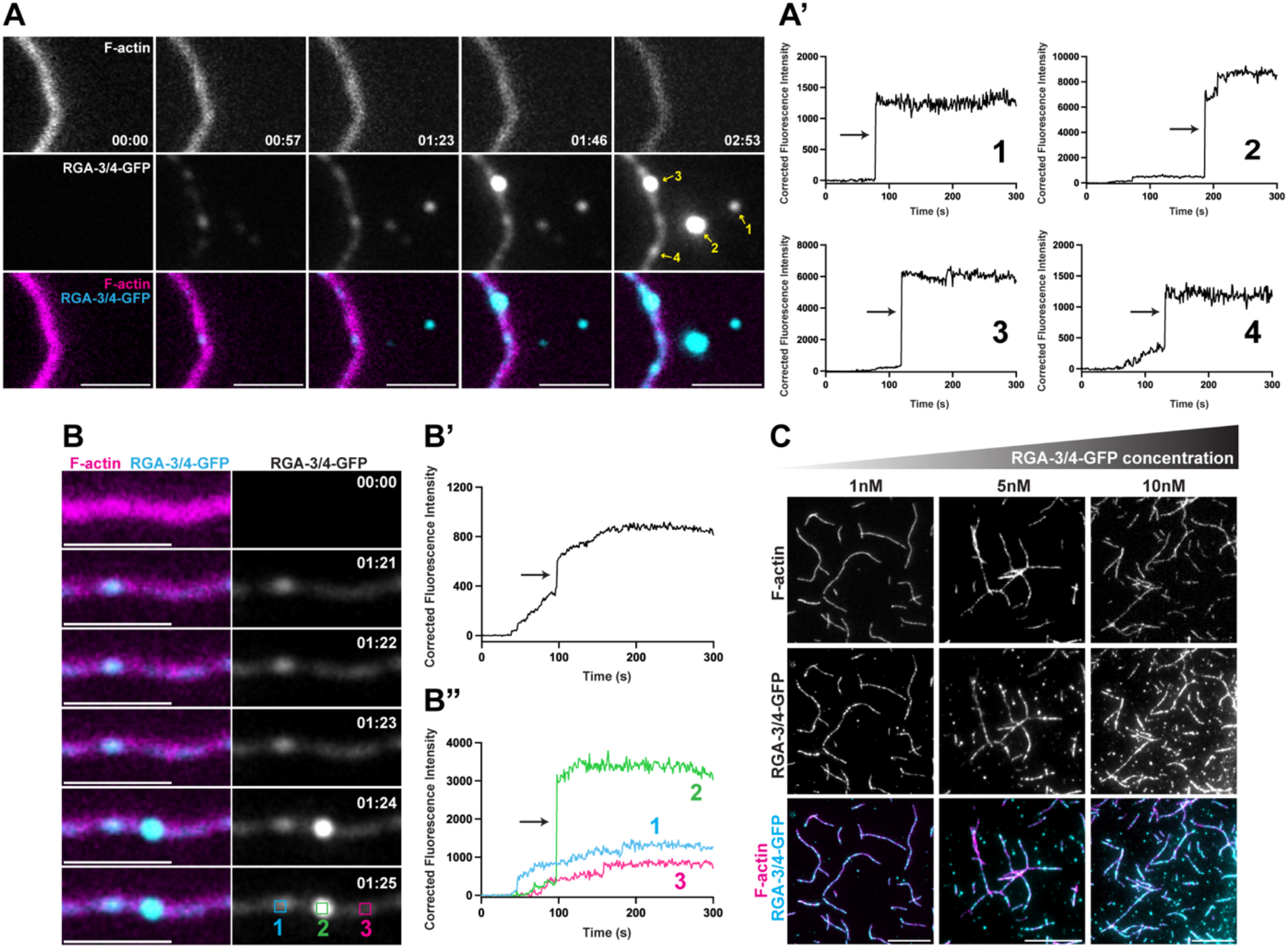
RGA-3/4-GFP clusters abruptly land on F-actin and background regions. (A) Time-lapse images showing RGA-3/4-GFP clusters on F-actin or background regions. Yellow arrows indicate clusters analyzed for (A’) corrected fluorescence intensity over time. Black arrows indicate abrupt landing of RGA-3/4-GFP clusters. (B) Time-course montage of RGA-3/4-GFP cluster abruptly landing on F-actin. Corrected fluorescence intensity over time of RGA-3/4-GFP measured along (B’) whole filament and (B”) subregions of the filament. (C) Representative final images of RGA-3/4-GFP with F-actin at increasing RGA-3/4-GFP concentration. Scale bars: 2*μ*m (A, B) and 10*μ*m (C).

**Supplemental Figure S4.**
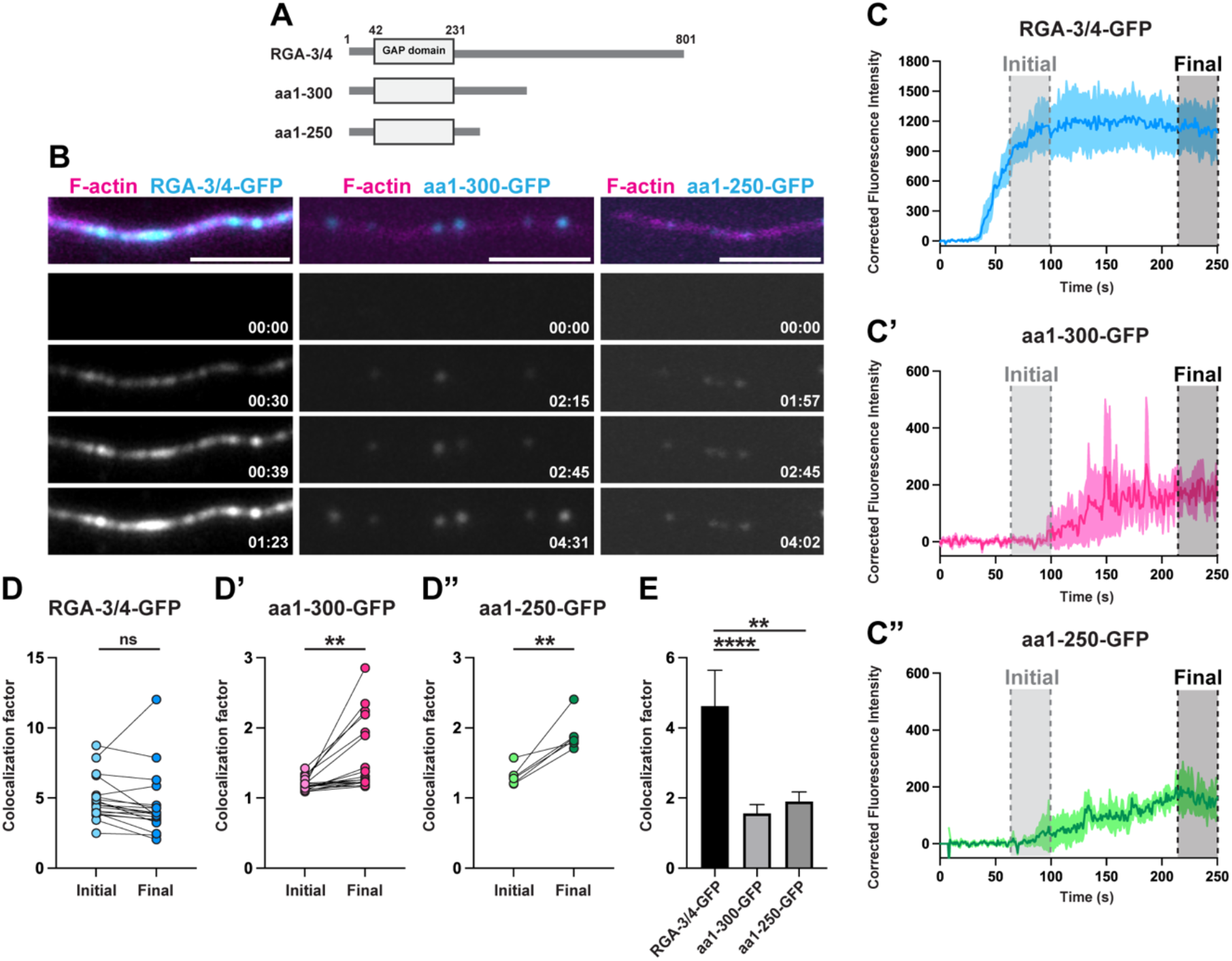
aa1-300 and aa1-250 exhibit delayed and reduced localization on individual actin filament. (A) Schematic diagrams of wildtype and truncated RGA-3/4 constructs. (B) Time-course montage of wildtype or truncated RGA-3/4-GFP proteins accumulating on F-actin. Scale bar: 3*μ*m. (C-C”) Corrected fluorescence intensity over time of wildtype or truncated RGA-3/4-GFP proteins measured along subregions of the filament. Mean±SD is shown. (D-D”) Quantification of colocalization between F-actin and wildtype or truncated RGA-3/4-GFP proteins at initial and final time points. Measurements from 2 experiments. **=0.0023 (aa1-300-GFP); **=0.0036 (aa1-250-GFP). Paired t-test was performed for statistical analysis. (E) Comparison of colocalization factor between wildtype and truncated RGA-3/4-GFP proteins at final time points. Measurements from 2 experiments. **=0.0014; ****<0.0001. One-way ANOVA with Tukey post-hoc test for multiple comparisons was performed for statistical analysis.

